# Biophysical insights into a highly selective L-arginine-binding lipoprotein of a pathogenic treponeme

**DOI:** 10.1101/350942

**Authors:** Ranjit K. Deka, Wei Z. Liu, Shih-Chia Tso, Michael V. Norgard, Chad A. Brautigam

## Abstract

Biophysical and biochemical studies on the lipoproteins and other periplasmic proteins from the spirochetal species *Treponema pallidum* have yielded numerous insights into the functioning of the organism’s peculiar membrane organization, its nutritional requirements, and intermediary metabolism. However, not all *T. pallidum* proteins have proven to be amenable to biophysical studies. One such recalcitrant protein is Tp0309, a putative polar-amino-acid-binding protein of an ABC transporter system. To gain further information on its possible function, a homolog of the protein from the related species *T. vincentii* was used as a surrogate. This protein, Tv2483, was crystallized, resulting in the determination of its crystal structure at a resolution of 1.75 Å. The protein has a typical fold for a ligand-binding protein, and a single molecule of L-arginine was bound between its two lobes. Differential scanning fluorimetry and isothermal titration calorimetry experiments confirmed that L-arginine bound to the protein with unusually high selectivity. However, further comparison to Tp0309 showed differences in key amino-acid-binding residues may impart an alternate specificity for the *T. pallidum* protein.

---

The acquisition of nutrients is a central requirement for all organisms, but one that is particularly nuanced in commensal, symbiotic, parasitic, and facultative pathogenic bacteria. These organisms must often contend with widely varying cycles of nutrient repletion and scarcity while simultaneously evading or subverting host immune surveillance. The evolutionary pressures on such bacteria are largely a product of their habitats. For example, a facultative pathogen likely harbors genes for use in its free-living state and also those activated when infecting a host. Whereas this strategy imparts maximum flexibility, there is a metabolic cost for maintaining genes that may or may not be utilized during an organism’s existence. On the other end of the spectrum, through evolution, an obligate parasitic bacterium may have jettisoned genes for certain catabolic pathways because it lives in an environment with ready supplies of given nutrients. The energetic advantage is countered by the fact that a paucity of the nutrient in the host cannot be easily countered and can lead to death of the organism.

Pathogenic bacteria often acquire essential metabolites from their environments using various transport systems to translocate the nutrient from the external milieu (i.e. the host) to the cytoplasm. One of the best characterized of such systems is the ABC transporters (1, 2). ABC transport systems typically are composed of multiple polypeptides: a ligand-binding protein (LBP) exists outside the cytoplasm (in the periplasm of Gram-negative bacteria or attached to the outside of Gram-positive bacteria), tasked with binding the target nutrient; a permease that spans the cytoplasmic membrane and receives the nutrient from the LBP; and a cytoplasmic ATP-binding protein, which provides the energy for transporting the nutrient against a free-energy gradient by hydrolyzing ATP. ABC transporters are known for a very wide variety of ligands (2–4), including sugars, amino acids, metal ions, and polyamines.

*Treponema pallidum*, the obligate human spirochetal pathogen that causes syphilis, has a highly parsimonious genome, and thus lacks enzymes for the *de novo* synthesis of many metabolites, including most amino acids (5). In an ongoing effort to understand the metabolic requirements of *T. pallidum*, we became interested in the gene product of *tp0309*, which putatively encodes an LBP for a polar amino-acid transporter. Experience has shown that these annotations must be experimentally verified (6, 7), as they can be inaccurate. Efforts to produce Tp0309 for biophysical and biochemical characterization were hampered by insolubility of the protein (despite the deployment of multiple solubilization and hyper-expression strategies). However, we noticed that the Tv2483 protein of the oral pathogen *Treponema vincentii* qualified as a homolog of Tp0309. We reasoned that characterization of this protein may further the understanding of Tp0309, and, ultimately, the nutritional requirements of *T. pallidum*.

To this end, we hyper-expressed Tv2483 in *E. coli* and determined its crystal structure. Its fold and the discovery of a single molecule of L-arginine bound to the protein prompted us to hypothesize that Tv2483 is an L-arginine-binding LBP. After a sequence of solution biophysics studies and structural comparisons to other L-arginine-binding LBPs, a view arose of Tv2483 as a highly selective LBP whose only cognate amino acid is L-arginine. These studies offered new insights into whether Tp0309 also fulfills this function.

## Results

### Preparations of Tv2483 are monodisperse and yield monomeric protein in solution

Most LBPs are monomers in solution (4). However, there are a few LBPs that have higher-order oligomeric states. These include LBPs for tripartite ATP-independent periplasmic transporters (8, 9) and for an ABC transporter for L-arginine (10). We therefore performed solution studies to establish the oligomeric status of Tv2483.

Purified Tv2483 (see Experimental Procedures) was subjected to dynamic light scattering (DLS) to assess the size and monodispersity of the preparation. The resulting autocorrelation function could be fitted with a cumulant analysis (11, 12), resulting in a polydispersity index of 0.09 ± 0.05. Fitting the data to a regularized distribution of hydrodynamic radii (*R*_*H*_) demonstrated that the vast majority (96 ± 2%) of the scattered-light intensity could be accounted for by a single species with an intensity-weighted *R*_*H*_ of 2.87 ± 0.16 nm (Fig. 1). Another species was usually observed, but its size varied greatly (*R*_*H*_ = 60 ± 30 nm); the wide variation was likely due to the small signal for this species compared to the noise value. These measurements were consistent with the notion that the Tv2483 preparation was dominated by a monomeric form of the protein.

**Figure 1.**
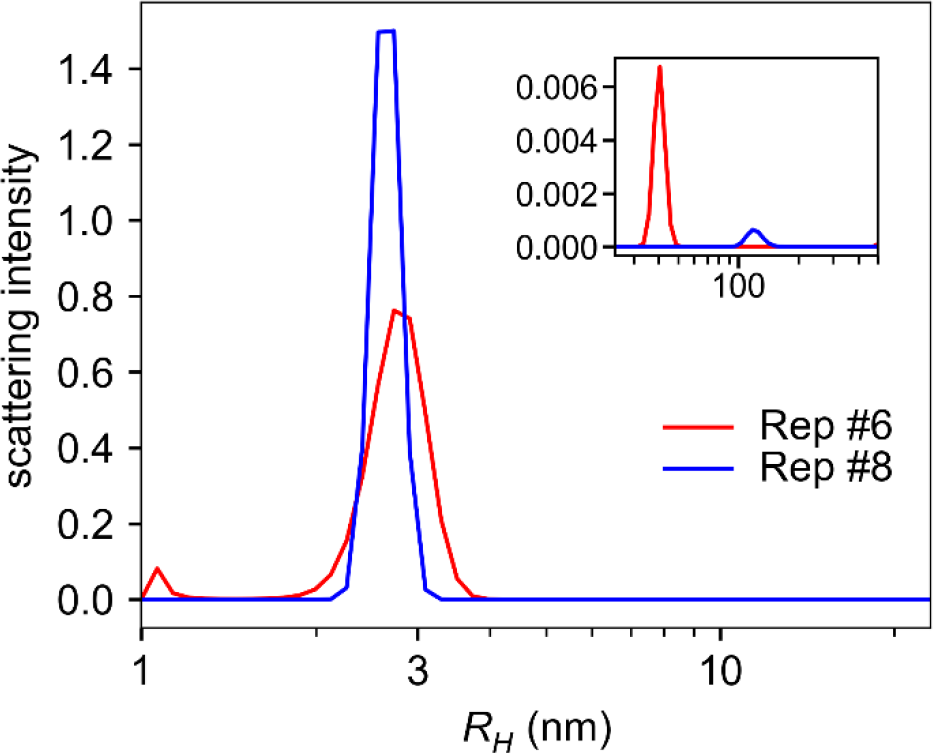
Hydrodynamic radius distributions of purified Tv2483. Two of the nine replicates (see Experimental Procedures) are shown, with coloration as described in the inset legend. The inset graph shows peaks at higher *R*_*H*_ values; note the scaling of the y-axis of this inset, as these are minority species.

To help to confirm this hypothesis, we conducted analytical ultracentrifugation (AUC) experiments on Tv2483 in the sedimentation velocity (SV) mode. Three separate experiments at different concentrations (23, 7.3, and 2.3 μM) were performed. Using the *c*(*s*) methodology to analyze these data, we observed a single peak in the distributions, representing an average molar mass of 30,500 ± 600 g/mol in the main peak (at an average experimental *s*-value of 2.429 ± 0.007 S) of the resulting distribution (Fig. 2). When compared to the known molar mass of the protein construct (30,165 g/mol), these SV experiments buttress the notion that Tv2483 is a monomer in solution.

**Figure 2.**
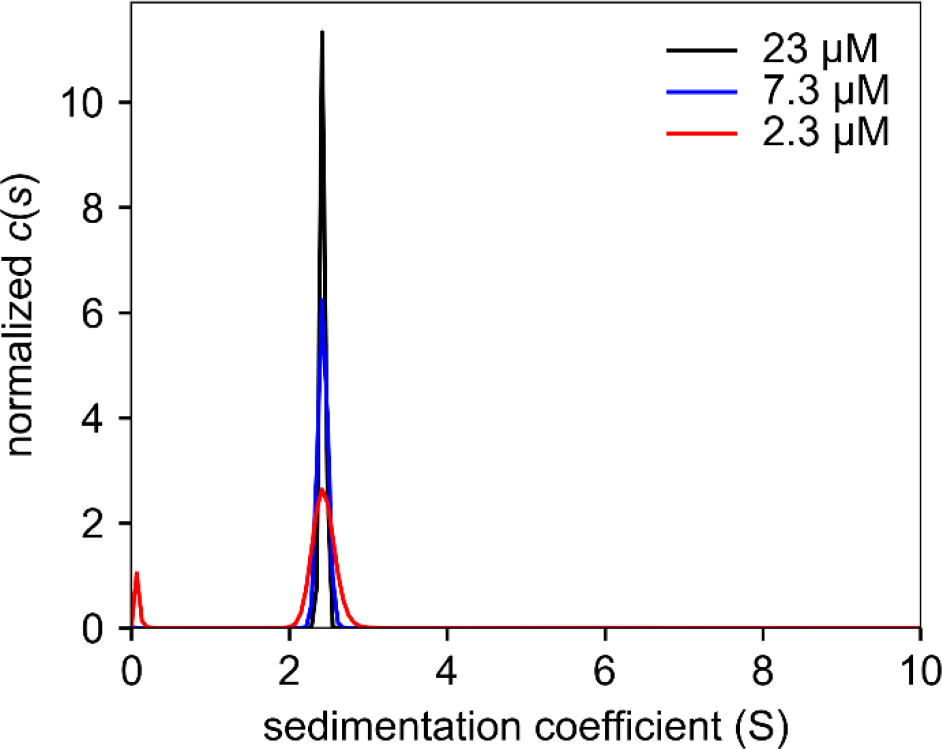
SV-derived *c*(*s*) distributions for purified Tv2483. The distributions have been normalized by the areas under the respective curves. The inset legend shows the concentrations used.

### The crystal structure of Tv2483 reveals an L-arginine-binding site

Tv2483 crystallized readily, and the crystals diffracted X-rays to a resolution of 1.75 Å (Table I). The crystal structure was determined using molecular replacement (see Experimental Procedures), revealing a single monomer of the protein in the asymmetric unit. The overall structure of the protein (Fig. 3) was bilobate; in keeping with the nomenclature for other amino-acid-binding proteins, we called these subdomains “Lobe I” and “Lobe II”. Lobe I (residues 4-99; 200-246) consisted of a central, five-stranded β-sheet, and the order of strands in the sheet was β2-β1-β3-β5-β4 (numbered according to order in the primary structure); all of the strands were parallel except β5. There was one α-helix near to the amino-terminus of the protein that formed a “cap” on Lobe I; most other helices were packed onto the faces of the sheet. Lobe II (residues 104-194) also comprised a single β-sheet flanked by helices. The strand order in this case was β3-β2-β4-β1-β5, with, β1 being the anti-parallel strand. Again, α-helices stacked on the faces of the sheet. Connecting the two lobes were two regions (residues 100-103 and 195-199) that adopted the secondary structure of β-strands.

**Figure 3.**
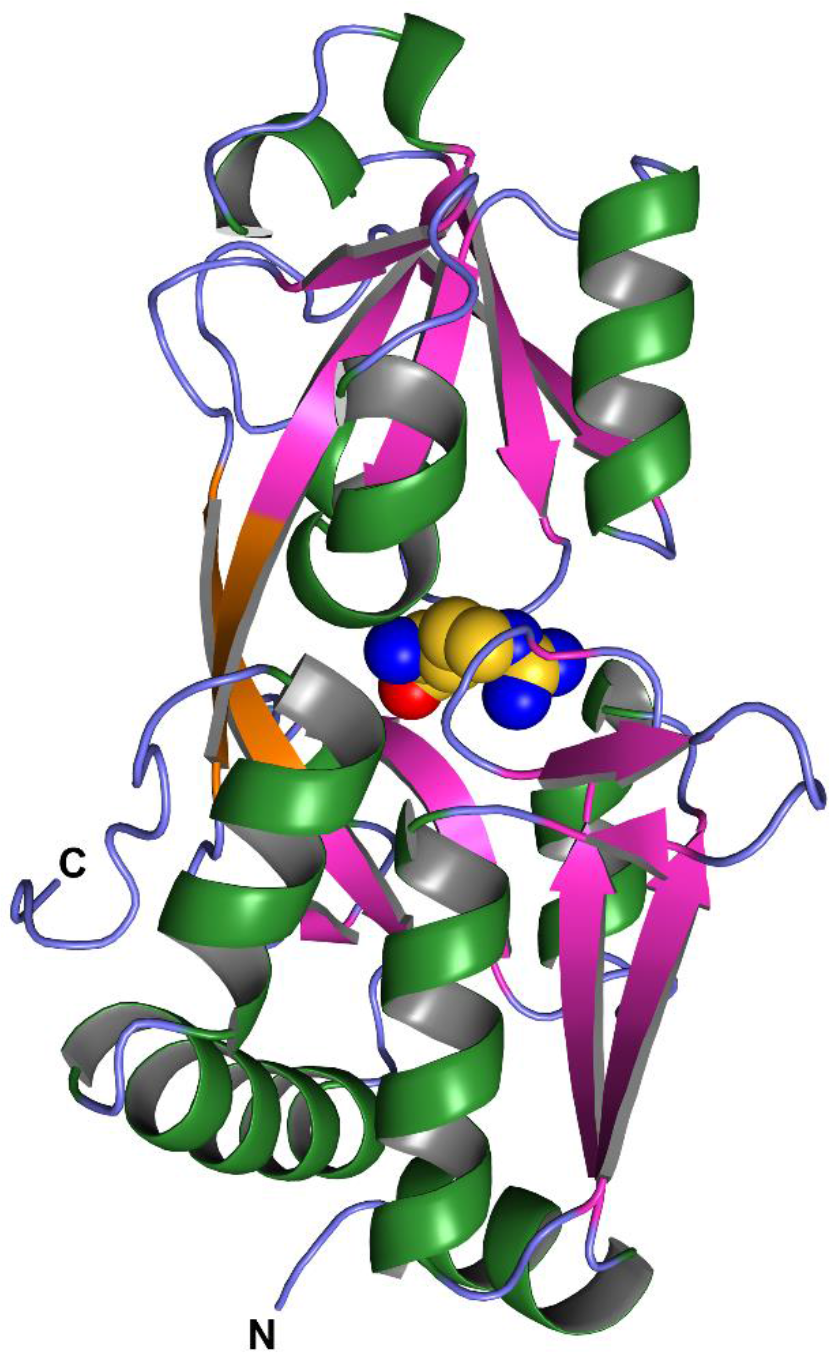
Overall crystal structure of Tv2483. The α-helices are colored green, the β strands
purple, and the linker region orange. Regions that are visible but without regular secondary structure are colored light blue. The amino-(N) and carboxyl-(C) termini are labeled. The bound molecule of L-arginine is shown as spheres, with carbon atoms colored yellow, nitrogens blue, and oxygens red.

**Table I.** Data collection, phasing, and refinement statistics.

| **Data Set** | Crystal I | Crystal II |
| --- | --- | --- |
| PDB Accession No. | N/A | 6DET |
| **Data Collection** |  |  |
| Space Group | C222 | C222 |
| Unit Cell Dimensions (Å) |  |  |
| a | 105.0 | 104.6 |
| b | 131.1 | 130.4 |
| c | 48.2 | 49.0 |
| $\alpha(^{\circ})$ | 90 | 90 |
| $\beta(^{\circ})$ | 90 | 90 |
| $\gamma(^{\circ})$ | 90 | 90 |
| Resolution (Å) | 48.2-1.90 (1.93-1.90)<sup>a</sup> | 81.60-1.75 (1.78-1.75) |
| Completeness (%) | 99.3 (93.9) | 97.8 (98.2) |
| Multiplicity | 8.0 (5.9) | 4.4 (4.5) |
| Unique Reflections | 26,530 (1,227) | 33,458 (1,639) |
| $R_{merge}^b$ | 0.087 (0.826) | 0.059 (0.742) |
| $\langle I \rangle / \sigma_I$ | 26.4 (2.0) | 22.4 (2.0) |
| Wilson B (Å<sup>2</sup>) | 24.5 | 27.0 |
| **Refinement** |  |  |
| Resolution (Å) |  | 42.0-1.75 |
| No. Residues |  | 243 |
| No. Non-Solvent Atoms |  | 1,944 |
| No. Solvent Atoms |  | 100 |
| Maximum-Likelihood |  |  |
| Coordinate Error (Å) |  | 0.17 |
| **Average B-factors** |  |  |
| Non-Solvent (Å<sup>2</sup>) |  | 42.1 |
| Ligand (Å<sup>2</sup>) |  | 32.7 |
| Solvent (Å<sup>2</sup>) |  | 41.6 |
| **R-values** |  |  |
| $R_{work}^c$ | | 0.177 |

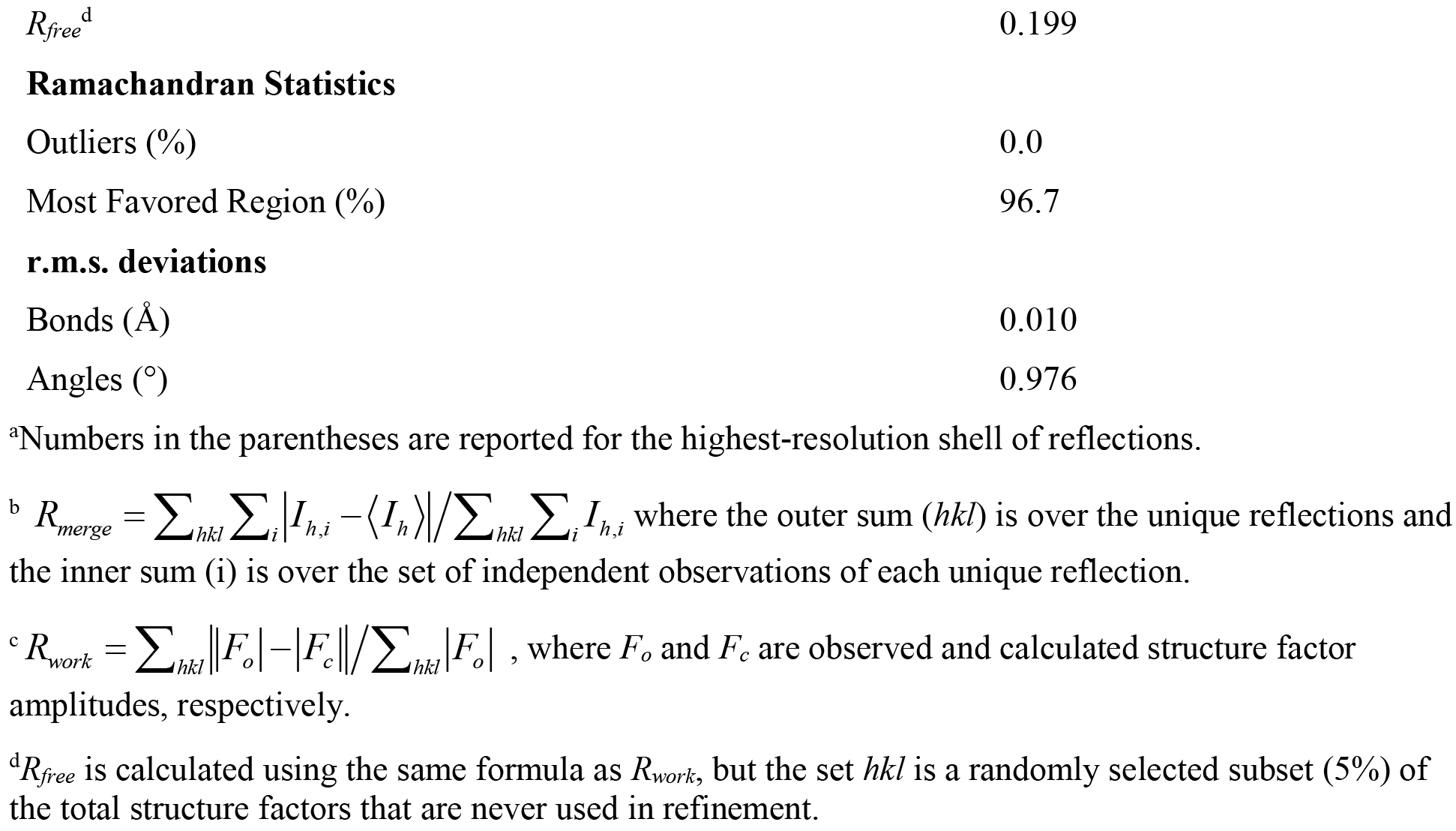

The structural features described above have been observed in other LBPs. Indeed, the overall topology, including the two connecting regions between the lobes, placed the protein in “Class II” of the classification scheme of Fukami-Kobayashi et al. (13). The protein was a member of Cluster F in the more detailed taxonomy put forth by Berntsson et al. (14). Notably, Cluster F proteins usually have linker regions devoid of regular secondary structure, whereas those in Tv2483 clearly are β-strands that are hydrogen-bonded to each other (Fig. 3).

Between the two lobes was clear electron density for a single molecule of L-arginine (Fig. 4A). The amino acid engaged the both protein lobes via a network of hydrogen bonds and putative salt bridges (Fig. 4). These interactions could be grouped into three categories: (i) hydrogen bonds/putative salt bridges from protein side chains, (ii) hydrogen bonds from the protein’s main chain, and (iii) hydrophobic or stacking interactions. In the first group, the amino-and carboxylate moieties of L-arginine were contacted by the presumably charged side chains of D173 and R89, respectively; the latter interaction was bidentate. Another bidentate interaction was between the guanidinium group of the L-arginine and the carboxylate moiety of E128. The nitrogen from the indole group of W64 formed a putative hydrogen bond with the carboxylate of the L-arginine, and the side-chains of two serine residues (S23 and S84) also interacted with the amino acid. In the second group, main-chain oxygen atoms of S81 and G82 apparently formed hydrogen bonds with the guanidinium group, and the main-chain nitrogen of S84 appeared to interact with the carboxylate of the L-arginine. The third group of contacts featured the guanidinium group and methylene moieties of the L-arginine sandwiched between the hydrophobic side chain of W64 on one side and those of V26 and Y176 on the other.

**Figure 4.**
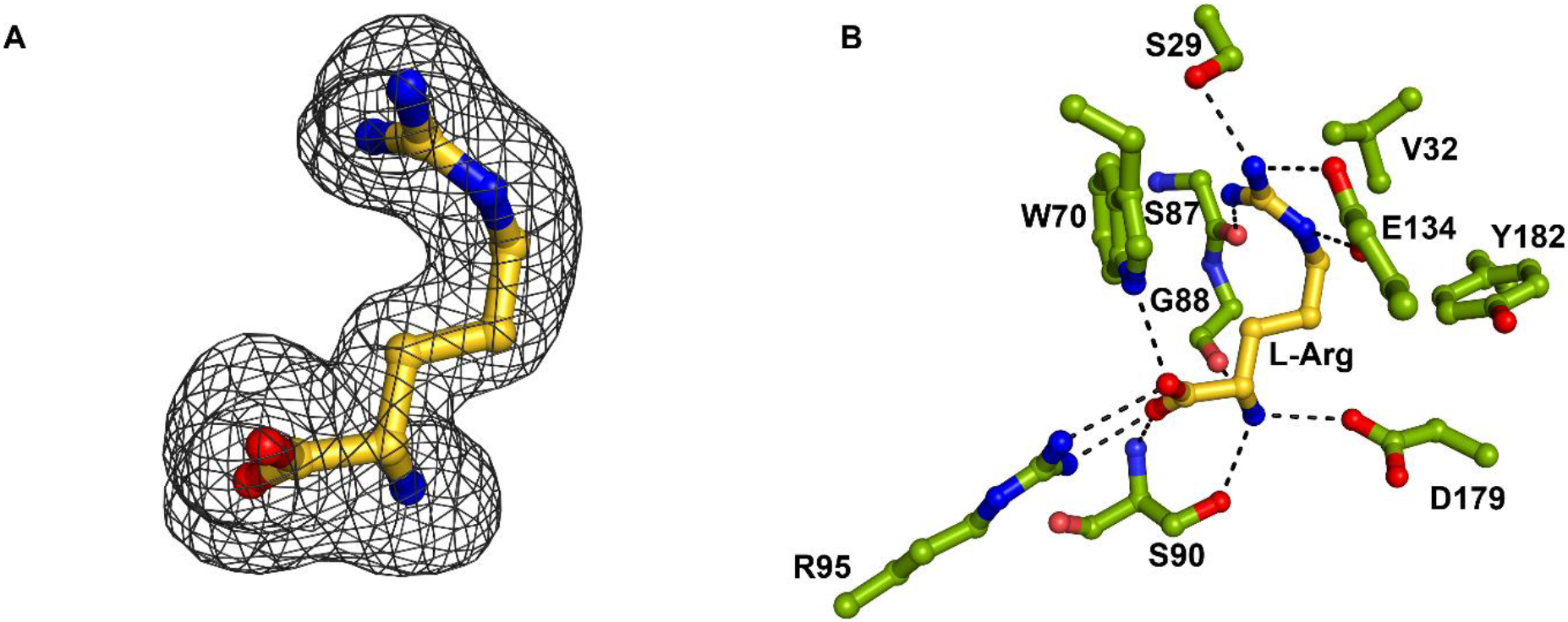
Details of the interaction between Tv2483 and L-arginine. (A) Electron density for the bound L-arginine. A kicked *mF*_*o*_ - *DF*_*c*_ omit map contoured at the 3-*σ* level is shown superposed on the final, refined position of the L-arginine bound to Tv2483. Colors are those established in Fig. 3. (B) Contacts between Tv2483 and the bound L-arginine. Putative hydrogen bonds or salt bridges are shown as dashed black lines. The L-arginine is colored as in Fig. 3, and the color scheme is the same for protein-derived residues except for the carbon atoms, which are shown in green. The side chain of S81 was omitted for clarity.

### Tv2483 is an L-arginine-specific binding protein

Because many basic-amino-acid-binding proteins have multiple specificities (15, 16), we initiated studies to determine which amino acids bound to Tv2483. We first used differential scanning fluorimetry (DSF) (17) to screen a panel of amino acids and other nitrogen sources for molecules besides L-arginine that could bind to Tv2483 (Fig. 5). Although L-arginine did clearly appear as a positive “hit” in the assay (shifting the *Tm* of the protein by +10 °C), the only other amino acid that had any effect was a nearly isosteric analog of L-arginine, L-citrulline.

**Figure 5.**
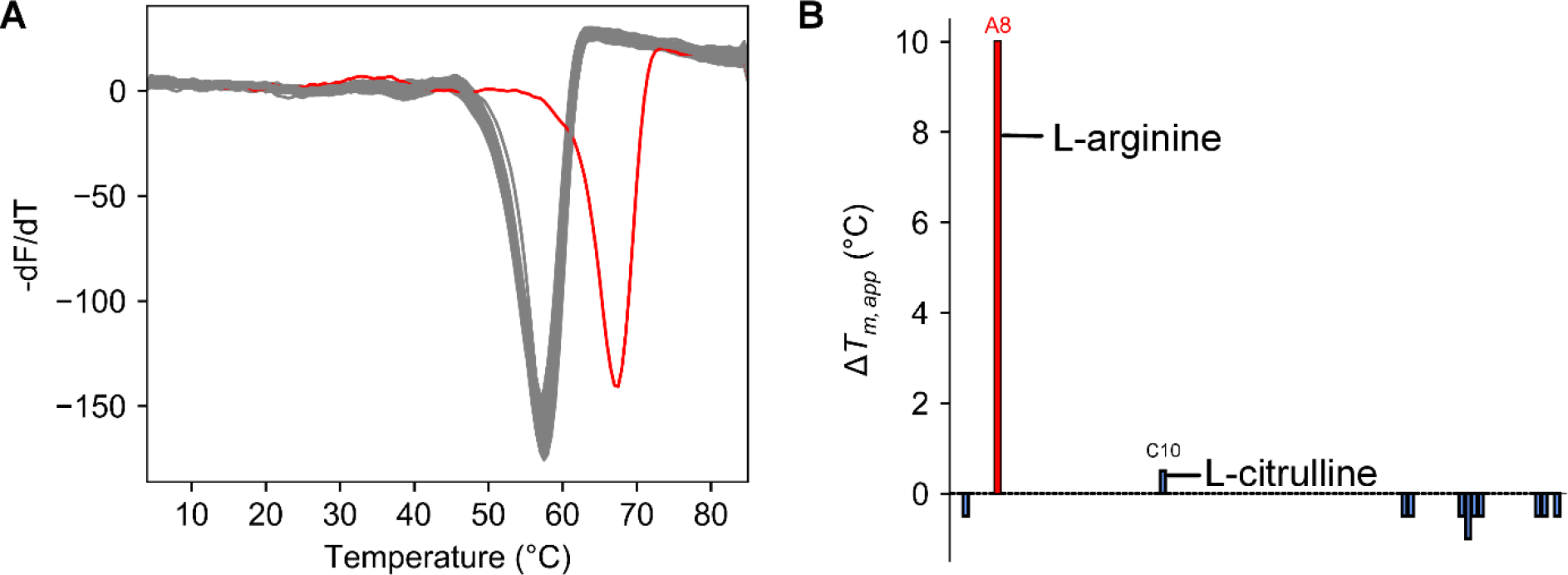
DSF of Tv2483 in the presence of nitrogen sources. (A) The –*dF/dT* traces. The red trace is that of well A8, which contains L-arginine; the gray lines are the traces for all other wells. (B) *T*_*m*_ shifts. On the *x-*axis is the well identity, with the size of the bar indicating the magnitude of the shift in *T*_*m*_ of the respective well compared to the water control. Well A8 is depicted in red. The wells containing L-arginine (A8) and L-citrulline (C10) are labeled.

Isothermal titration calorimetry (ITC) was employed to quantify the affinity of L-arginine for Tv2483. Our initial efforts were thwarted by a lack of a heat signal in the resulting thermograms (not shown). We surmised that L-arginine was co-purifying with the protein, preventing a discernible signal from appearing when titrating in more L-arginine. We attempted to remove the amino acid by a denaturation/renaturation scheme (6) and by extensive dialysis, but neither approach succeeded. Taking a cue from the DSF assay, we devised a method using high concentrations of L-citrulline to compete for binding to the protein, followed by removal of the competitor using repeated cycles of dilution and concentration. We estimated that this strategy rendered approximately 65% of the protein competent to bind to L-arginine (see Experimental Procedures).

L-arginine bound avidly to L-citrulline-treated Tv2483 (Fig. 6A; Table II). Indeed, the isotherms derived from titrations of L-arginine into Tv2483 could not be analyzed to arrive at an accurate *K*_D_ because the binding was so avid. Instead, we incorporated these isotherms into a global analysis that included titrations of L-citrulline into Tv2483 and L-arginine into a mixture of L-citrulline and Tv2483 (18, 19). This analysis allowed us to determine that the protein’s *K*_D_ for L-arginine was 6 [2, 12] nM (we include the 68.3% confidence intervals in brackets in this work). Another result from this global analysis was the determination of the *K*_D_ for L-citrulline, which was 11.4 [10.0, 13.0] μM, i.e. almost 2,000-fold that for L-arginine.

**Figure 6.**
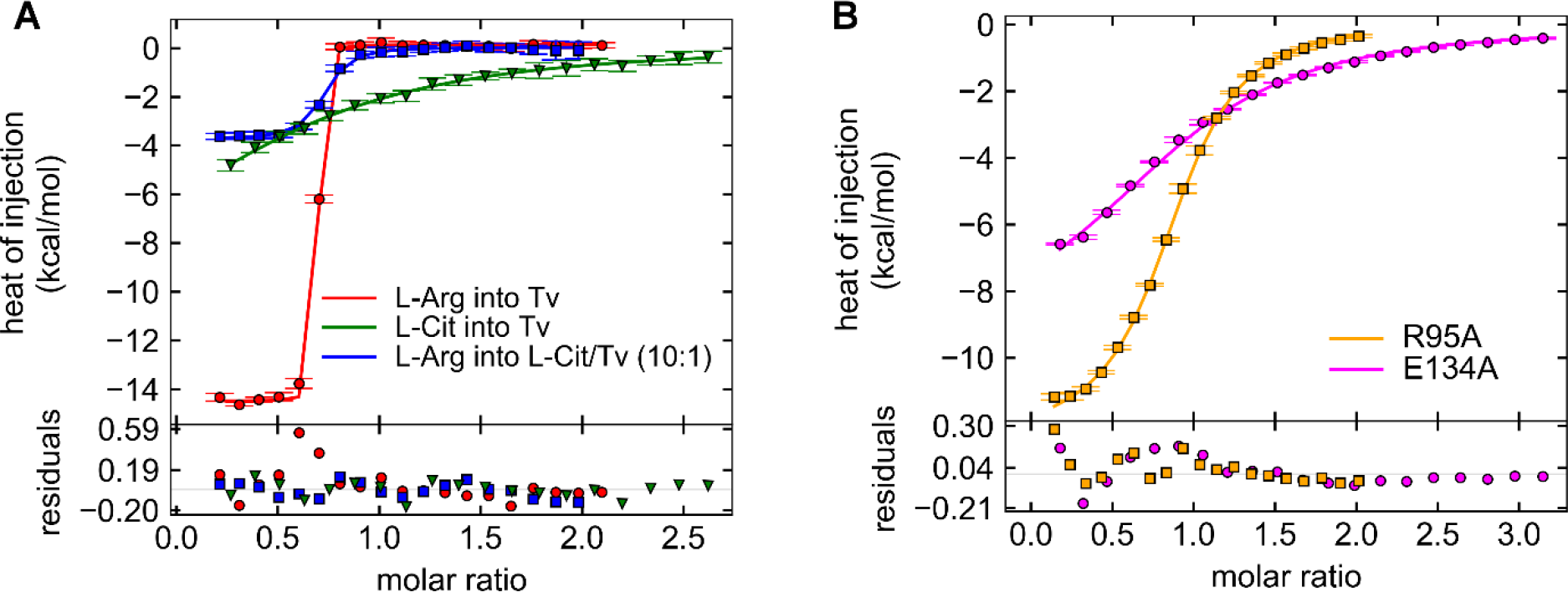
ITC of Tv2483 binding to L-arginine. (A) Isotherms for the wild-type protein. Three isotherms of the five used in the global analysis are shown, colored according to the inset legend. In the upper graph are the isotherms, with the integrated heats depicted as markers and the results of the global fit to the data shown as lines. Residuals between the data and the fit lines are shown in the lower plot. “L-Arg” is L-arginine, “L-Cit” is L-citrulline, and “Tv” is Tv2483. (B) Isotherms for L-arginine titrated into the mutant proteins. Conventions established in part (A) are followed here.

**Table II.** *K*_D_s of L-arginine binding to Tv2483 and its variants.

| **Protein** | **Ligand** | **$K_D$ (μM)** |
| --- | --- | --- |
| wild-type | L-arginine | 0.006 [0.002, 0.012]<sup>a</sup> |
| wild-type | L-citrulline | 11.4 [10.0, 13.0] |
| wild-type | L-lysine | N.H.<sup>b</sup> |
| wild-type | L-ornithine | N.H. |
| R95A | L-arginine | 6.3 [5.9, 6.8] |
| E134A | L-arginine | 31 [26, 37] |

<sup>a</sup>68.3% confidence intervals determined by error-surface projection (52) are shown in square brackets.
<sup>b</sup>N.H.: no heat of binding was observed during the titration.

We also studied the binding of L-lysine and L-ornithine to Tv2483. Using Tv2483 that had been L-citrulline treated, there was no heat evident upon titration of neither of these basic amino acids (not shown). Thus, the results we achieved with ITC were entirely consistent with the DSF assay, and strongly suggested that Tv2483 is an L-arginine-specific binding protein.

To validate the identities of the amino-acid residues that we observed in the Tv2483 crystal structure, we mutated two of the contacting residues to alanine and studied their binding to L-arginine using ITC (Fig. 6B; Table II). Notably, we found that no L-citrulline treatment was necessary for the mutant proteins; ostensibly, the weakened binding characteristics of the mutated proteins minimized the amount of co-purified amino acids in these preparations. The first mutation was targeted at the putative salt bridge between R89 and the carboxylate moiety of the bound L-arginine. Substitution of this residue with alanine raised the *K*_D_ by about 1,000X (6.3 [5.9, 6.8] μM). The second mutation was changing to alanine the glutamate whose side chain made a presumed salt bridge with the guanidinium moiety of the bound amino acid (E128). This mutation was more deleterious to L-arginine binding than the previous, displaying a *K*_D_ that was higher than the wild-type by approximately 5,000X (31 [26, 37] μM). These results allowed us to conclude that the binding site observed in the crystal structure is the operative one in solution, and that the abrogation of a single salt bridge, while not enough to fully eliminate binding, has significantly negative effects on the binding of L-arginine to Tv2483.

### Comparison of Tv2483 to other LBPs helps to define the specificity determinants

Database searches (20) for similar tertiary structures revealed a large number of other amino-acid-binding LBPs. Among them are proteins that bind L-glutamate or L-histidine. However, we chose to focus on those that interact with basic amino acids, as does Tv2483 (see Table III).

**Table III.** Comparisons of Tv2483 to other L-arginine-binding LBPs.

| **Protein** | **r.m.s.d. ($\text{\AA}$)** | **No. comparable $C_\alpha$ atoms** |
| --- | --- | --- |
| StLAO | 2.0 | 227 |
| GsArtJ | 2.1 | 229 |
| STM4351 | 1.9 | 218 |

The basic-amino-acid-binding LBPs identified in the searches had a wide range of specificities and affinities for their respective ligands. For example, several structures of lysine-arginine-ornithine (LAO) binding proteins were returned. The best characterized of these is the LAO protein from *Salmonella typhimurium*, which we term “StLAO” henceforth. The root-mean-squared deviation (r.m.s.d.) in the positions of the superposed C_α_ atoms was 2.0 Å over 227 comparable atoms. StLAO had no appreciable specificity among its three eponymous amino acids, with *K*_D_ of 15, 14, and 29 nM for the three, respectively (15). Like Tv2483, StLAO sandwiched the side chain of the bound L-arginine between two hydrophobic side chains (Y14 and F52), and there were numerous apparent hydrogen bonds between the protein and the bound amino acid (Fig. 7A). In StLAO, none of the interactions between the protein and the guanidinium group of the bound L-arginine were bidentate. Notably, the bound amino acid is in a different conformation in Tv2483 than that displayed in the binding site of StLAO; the L-arginine side chain is “bent” in former structure relative to the “straight” conformation observed in the latter. Indeed, it is this bend that allows E128 to engage in its bidentate contact with the guanidinium group. In StLAO, the analogous residue is a leucine (L117), which has a van der Waals contact with the bound amino acid. Additionally (and not shown in Fig. 7A for clarity), a water molecule makes contact with the guanidinium group of the bound L-arginine.

**Figure 7.**
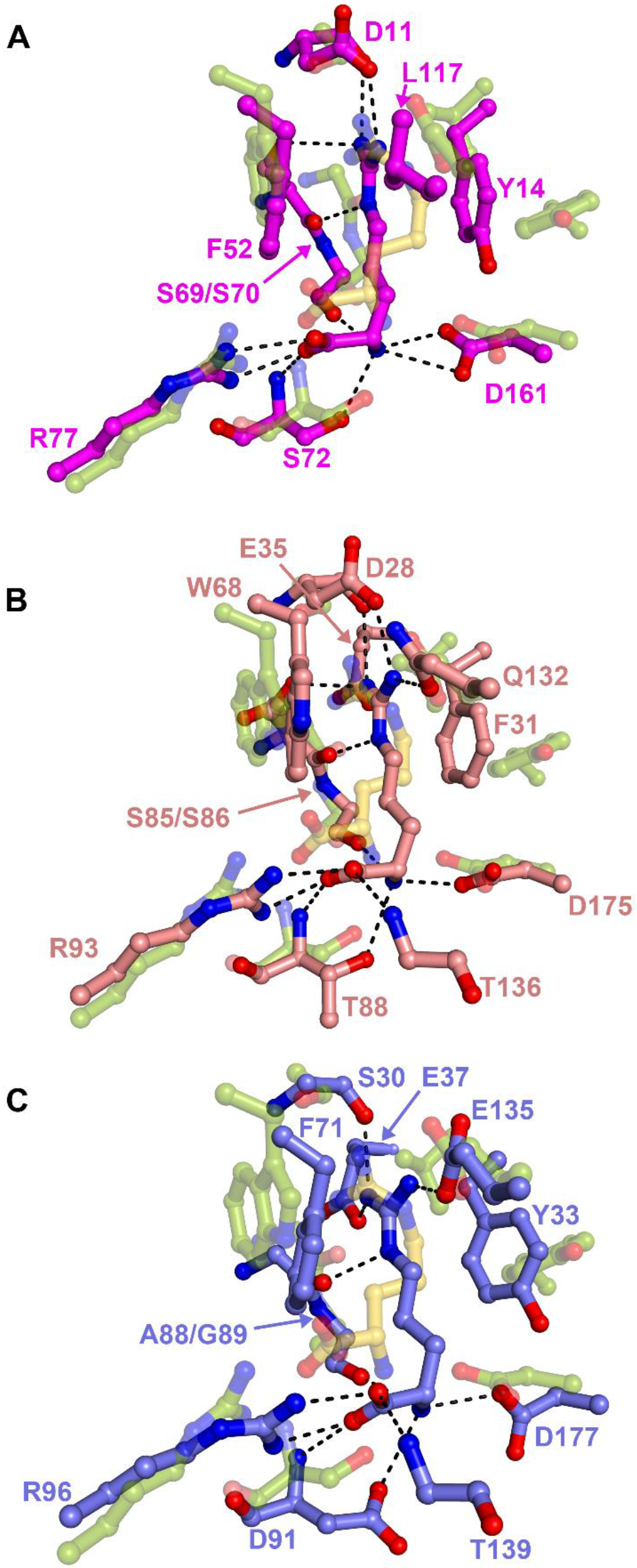
Superpositions of Tv2483 and other L-arginine-binding LBPs. Comparisons are shown for StLAO (A, magenta carbon atoms), GsArtJ (B, light red carbon atoms), and STM4351 (C, light blue carbon atoms). The residues presented in Fig. 4B for Tv2483 are shown semi-transparently here, in the color scheme of that figure. Some side chains and waters have been omitted for clarity. For STM4351, we used the amino-acid numbering in the PDB deposition, not that occurring in the attendant publication (21).

Another protein identified by our searches was ArtJ from *Geobacillus stearothermophilus* (16) (hereafter referred to as GsArtJ). The superposition of GsArtJ and Tv2483 resulted in an r.m.s.d. of 2.1 Å over 229 comparable C_α_ atoms. The overall themes of L-arginine binding displayed in StLAO were recapitulated in GsArtJ (Fig. 7B): the straight side chain of the bound amino acid was sandwiched between the side chains two hydrophobic residues (F31 and W68), the amino-and carboxylate moieties were contacted by polar or charged side chains, and the guanidinium group interacted with the side chains of several more polar/charged residues. Compared with the other L-arginine-binding proteins surveyed in this work, GsArtJ binds tightly to L-arginine, with a reported *K*_D_ of 40 nM (16). GsArtJ is also somewhat promiscuous, binding L-lysine and L-histidine in addition to L-arginine, and it appeared to use water molecules to aid in this binding plasticity, like StLAO does.

The final comparison presented in this report was between Tv2483 and STM4351 from *Salmonella enterica* (21) which, when superposed with Tv2483, exhibited a 1.9 Å r.m.s.d. over 218 aligned C_α_ atoms. This protein, though exhibiting a high degree of structural homology to StLAO, specifically bound to L-arginine; binding could only be detected to that amino acid and to *N*-methyl-L-arginine. The modest affinity of STM4351 for L-arginine (2.9 μM) indicates that the protein has a several-hundredfold higher *K*_D_ when compared to those of Tv2483, StLAO, or GsArtJ. The structure of STM4351 showed that the bound amino acid was recognized using contacts similar to those observed in Tv2483, StLAO, and ArtJ (Fig. 7C): charge-charge interactions at the amino-and carboxylate moieties of the bound amino acid, sandwiching of the L-arginine’s side chain between two hydrophobic residues, and numerous contacts to the bound guanidinium group. Notably, the side chain of the bound L-arginine was also in the straight conformation, as it was in both non-treponemal proteins discussed above. The changes that rendered STM4351 specific for L-arginine relative to StLAO focused on the replacement of a water molecule in the StLAO binding site with side chains making van der Waals contacts to the guanidinium group (21); waters in the binding site presumably contributed to the amino-acid promiscuity in StLAO and GsArtJ (16, 21, 22). Tv2483 employed a similar strategy, in that water was apparently excluded from the protein’s amino-acid binding site. The source of the relatively weak binding of L-arginine to StLAO was not evident from this comparison. A fourth L-arginine-binding protein identified in our structural-similarity searches, ArtJ from *Chlamydia pneumoniae* (23), was excluded from a more detailed analysis because of its relatively high r.m.s.d. when compared to Tv2483 (2.3 Å over 217 C_α_ atoms).

### Comparison of the Tv2483 and Tp0309 sequences reveals salient differences within the ligand-binding sites

Because Tv2483 is a homolog of Tp0309, we used a structure-based alignment algorithm (24) to examine whether Tp0309 was also likely to bind to L-arginine. *T. pallidum* has no enzymes for the *de novo* synthesis of L-arginine, and thus probably must import it by some means. Generally, the two primary structures aligned well, displaying an amino-acid identity of about 25% with few gaps and strongly corresponding secondary-structure predictions (Fig. 8). Two key L-arginine-binding residues were conserved between the two proteins: S23 and W64. However, the other residues were not strictly conserved; particularly problematic for the prospect of amino-acid binding by Tp0309 were the lack of charge conservation at the aspartic acid that bound the amino group (D173 corresponds to N200 in Tp0309) and the glutamic acid that interacted with the guanidinium group (the Tp0309 analog to E128 is H158). Also, the bidentate interaction observed in all L-arginine-binding proteins described above between (in Tv2483) R89 and the carboxylate of the bound amino acid was apparently not present in Tp0309 (it is H119 in that protein). These changes obscure the identity of the cognate ligand of Tp0309; although it is possible that the latter does bind L-arginine, that putative binding mode would differ from that observed for Tv2483 in this report. Regardless of these differences, the quality of the alignment implied that Tv2483 can serve as a good model for the overall fold of Tp0309.

**Figure 8.**
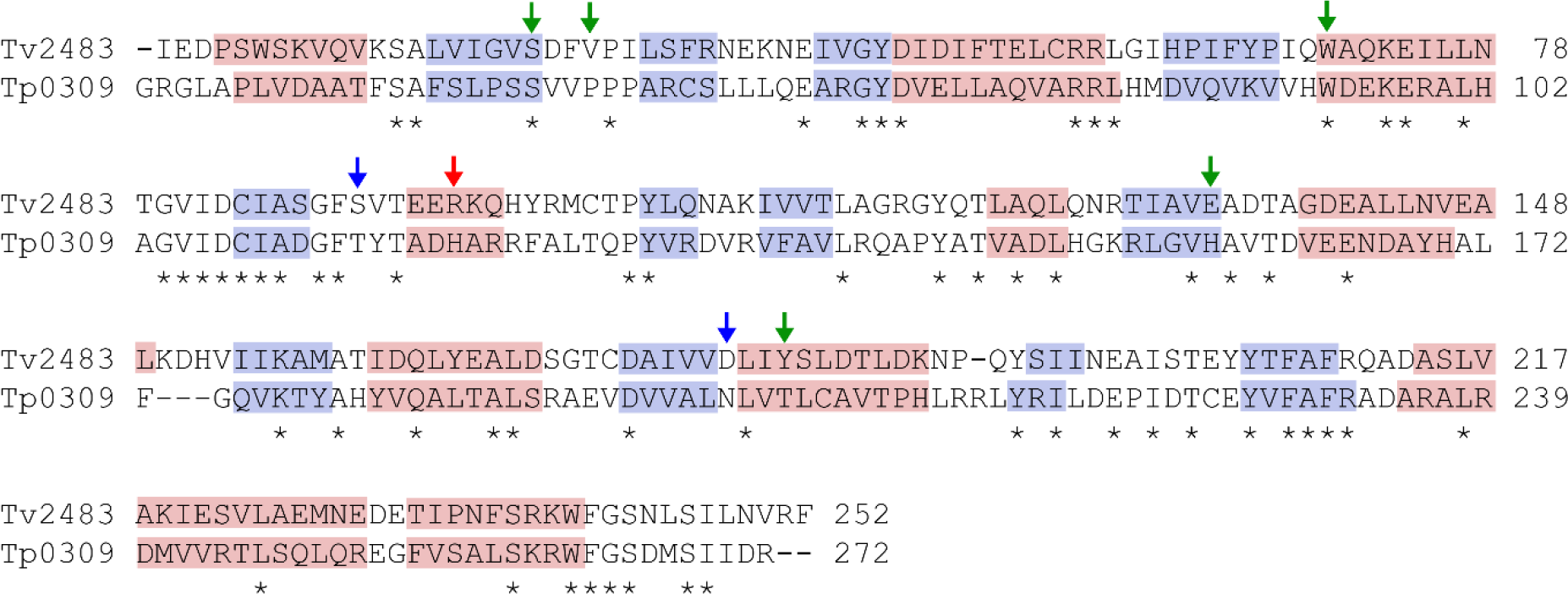
Sequence alignment of Tv2483 and Tp0309. The sequence numbering for Tp0309 is for the entire protein, as no signal sequences are evident near to its amino-terminus. Pink boxes indicate areas of predicted α-helices, whereas light-blue boxes depict predicted β-strands. Positions of amino-acid identities between the two proteins are marked with asterisks. Green arrows point out residues in Tv2483 whose side chains have contacts with the side chain of the bound L-arginine; blue arrows show those in contact with the amino group, and the red arrow shows the residue in contact with the carboxylate group.

## Discussion

Our results strongly support the contention that Tv2483 is a monomeric LBP that specifically and tightly binds to L-arginine (Figs. 1-3). Exclusion of water from the binding site and strong shape complementarity, first proposed to explain the specificity of STM4351 (21), appear to be operative in Tv2483 as well. The UniProt entry on Tv2483 notes that it has a periplasmic localization signal peptide at its amino-terminus; however, our investigations using the LipoP server (25) strongly suggested that it is a lipoprotein, like many LBPs of treponemes (e.g. (7, 26, 27)). Thus, the protein is probably anchored to the outer leaflet of the *T. vincentii* cytoplasmic membrane and binds L-arginine that enters the organism’s periplasm.

Many bacteria organize the genes necessary for a complete ABC transporter into a single operon. This fact conveniently enables the identification of an LBP’s cognate permease and ATP-hydrolyzing partners. This, however, is not the case for Tv2483; there are no nearby genes for other ABC transporter components except for Tv2484, which is apparently another LBP. This observation raises the possibility that the L-arginine import components are scattered in the *T. vincentii* genome or that Tv2483 (and Tv2484) dock on a permease that has a low binding-partner discrimination.

LBPs can be used for functions other than nutrient import in bacteria. For example, the MglB protein of *E. coli* can serve as both an LBP and a chemotactic sensor for D-galactose (28). Whereas it is tempting to speculate that the lack of co-operonic ABC components for Tv2483 could indicate a chemotactic function, the properties of the protein seem ill-suited for this purpose. Our experiences with the protein suggest that L-arginine is slow to dissociate from Tv2483 in the absence of an external impetus (see above and Experimental Procedures); this aspect of L-arginine binding does not comport with a chemotactic system that must be able to respond rapidly to changes in solute concentrations. Furthermore, the Tp0737 protein of *T. pallidum* offers an apparent precedent of a treponemal LBP that has no co-operonic ABC transporter apparatus (29).

A fundamental question raised by this study is how to rationalize the various binding affinities for L-arginine-binding LBPs. Tv2483, StLAO, and GsArtJ all bound the amino acid with *K*_D_s below 100 nM (Fig. 6 and (15, 16)). Given that the side chain of the bound L-arginine in only one of these structures is “bent” (that in Tv2483, Fig. 4), it was clearly indicated that this conformational variance does not manifest as a significant change in the free energy of amino-acid binding. However, the L-arginine-specific protein STM4351 has a much higher *K*_D_ than the other L-arginine-binding LBPs (2.9 μM), without an obvious loss in the number or quality of contacts to the bound amino acid. ITC of L-arginine binding to STM4351 (21) showed that the interaction was endothermic (having an observed Δ*H* of +1.4 kcal/mol), implying that binding was entropy-driven. Our studies of Tv2483 likely point to an exothermic heat of interaction for L-arginine binding (we observed a Δ*H* of −14.6 [−14.8, −14.5] kcal/mol; see Fig. 6A), although this cannot be definitively stated because of the ionization enthalpy of the buffer used in our studies (see Experimental Procedures and (30, 31)). If the observation of opposite binding enthalpies is accurate, these thermodynamic differences could reveal clues on the molecular mechanisms of tight versus modest L-arginine binding. Further calorimetric and structural studies would be necessary to explore this notion.

The linker region of Tv2483 comprised two β-strands (Fig. 3). LBPs of Cluster F (14) usually do not have regular secondary structure features in this region, as large interdomain motions, manifested as changes in backbone dihedral angles in the linker (32), are necessary. While GsArtJ also had β-strands in the linker, the length of the strands in Tv2483’s linker was exceptional. The implications of this observation for the range of motion of the two domains relative to one another are unknown at this time. Future efforts at obtaining an unliganded Tv2483 structure may reveal more about the plasticity (or lack thereof) in this region of the protein.

Our original homology search revealing that Tv2483 was a homolog of Tp0309 provided initial optimism that the *T. vincentii* homolog could serve as a surrogate in our efforts to characterize the intractable Tp0309 protein biochemically and biophysically. Although *T. vincentii* is not an obligate human pathogen like *T. pallidum*, it often is found in cases of periodontal disease, typically in association with other oral spirochetes and anaerobes (such as *Fusobacterium*) (33, 34). Thus, there are obvious salient biological differences between *T. pallidum* and *T. vincentii* and the human sites that they inhabit. Perhaps this biological difference, at least in part, explains the binding-pocket divergence that we now predict exists between Tv2483 and Tp0309, which could only be revealed by solving the crystal structure of Tv2483. Nonetheless, it remains possible that homologs of *T. pallidum* proteins found in other pathogenic spirochetes may continue to be useful for discerning biological and biochemical functions of other *T. pallidum* proteins.

## Experimental Procedures

### Protein Expression and Purification

To produce a recombinant derivative of TREVI0001_2483 (referred as “Tv2483”) in *Escherichia coli*, the DNA fragment encoding amino acid residues 25-272 of Tv2483 was PCR amplified from *Treponema vincentii* ATCC 700013 genomic DNA (ATCC, VA) by the polymerase incomplete primer extension (PIPE) cloning method using ends-specific primers (PIPE insert). The expression vector, pSpeedET (DNASU, Tempe, AZ), which encodes an N-terminal expression and purification hexa-histidine tag (MGSDKIHHHHHHENLYFQG), was PCR amplified with PIPE-vector primers. The PIPE-insert and PIPE-vector was mixed to anneal the amplified DNA fragments together (35). *E. coli* HK100 competent cells were transformed with the mixtures (PIPE-vector and insert) and selected for kanamycin resistance on LB agar plates. Cloning junctions/fragments were verified by DNA sequencing. A verified plasmid was then co-transformed with pGroESL (Takara USA, Mountain View, CA) into *E. coli* BL21 AI (Invitrogen, Waltham, MA) cells for soluble protein expression. *E. coli* BL21 AI cells were grown at 37 °C in LB medium containing 40 μg/mL of kanamycin and 30 μg/mL of chloramphenicol until the cell density reached an *A*_600_ of ~0.6. The cells were then induced for ~20 h with 0.2% (w/v) L-arabinose at 16 °C and harvested, and cell pellets were stored at −80 °C. The procedures for expression and purification of the recombinant proteins were essentially as previously described (36, 37).

### Site-Directed Mutagenesis and Protein Concentration Determination

For the construction of rTv2483 variants, the R89A and E128A mutations were individually introduced into the plasmid carrying the wild-type *tv2483* sequence using the QuikChange site-directed mutagenesis kit (Agilent Technologies). The mutation was confirmed by DNA sequencing. The mutant protein was expressed and purified as described above. Protein concentrations were determined in Sample Buffer (20 mM Hepes, 0.1 M NaCl, pH 7.5, 2 mM n-Octyl-β-D-glucopyranoside) using UV absorption at 280 nm. Extinction coefficients were calculated from the protein sequences using the ProtParam tool of ExPASy server (www.expasy.org).

### Protein Crystallization

Using the sitting-drop vapor-diffusion technique in 96-well plates with the PACT Crystallization Suite and Structure Screen 1 + 2 (Molecular Dimensions, Maumee, OH) and a crystallization robot (Crystal Gryphon, Art Robbins Instruments, Sunnyvale, CA), the initial crystallization conditions were determined at 20 °C. Crystallization screens for Tv2483 with a protein concentration of ~13 mg/ml (in Sample Buffer) yielded two conditions for diffraction-quality crystals (Crystal I and Crystal II). Crystal I appeared in Well A2 of the PACT screen (0.1 M SPG (succinic acid, sodium dihydrogen phosphate, and glycine) buffer, pH 5.0, 25% (v/v) PEG 1500) after 7 days. The crystals were transferred to the Stabilization Buffer I (0.1 M sodium citrate pH 5.0, 0.1 M NaCl, 25% PEG 1500, 5% (v/v) ethylene glycol), then serially transferred to solutions containing more ethylene glycol in increments of 10% to a final concentration of 25%. After about 1 min in this final mixture, the crystals were flash-cooled in liquid nitrogen. Crystal II appeared in well C7 of the Structure 1 + 2 screen (0.1 M HEPES pH 7.5, 10% (v/v) isopropanol, 20% (w/v) PEG4000. The crystal was moved into Stabilization Buffer II (0.1 M HEPES pH 7.5, 10% isopropanol, 20% PEG4000, 0.1 M NaCl, 2 mM n-Octyl-P-D-glucopyranoside, 5% (v/v) ethylene glycol), then transferred to Stabilization Buffer II supplemented to 15% ethylene glycol, where it resided for approximately 5 min before plunging into liquid nitrogen.

### X-ray Data Collection, Structure Determination, and Structure Refinement

X-ray diffraction data were collected at beamline 19-ID, the Structural Biology Center at Argonne National Laboratories. The crystals had the symmetry of space group C222 (Table I). An initial data set having a smallest *d*_*min*_ spacing of 1.9 Å was collected from Crystal I. The data were indexed, integrated, and scaled using HKL3000 (38). Hidden-Markov model searches (39) of the Protein Data Base showed that a histidine-binding protein, PDB accession number 4OHN, a histidine-binding protein from *Streptococcus pneumoniae* was the top match (there was a 25% sequence identity between the two). CLUSTALW (40) was used to align the sequences of 4OHN (no attendant publication) and Tv2483, and the molecular-replacement search model was constructed using Sculptor in the PHENIX program suite (41). Phaser (42) found one copy of the search model in the asymmetric unit (log-likelihood gain = 2,538), and the data and model were submitted to the ARP/wARP web server (43), which built 99% of the model. The remaining parts of the model were built in Coot (44). Rigid-body refinement, simulated annealing, individual B-factor, and positional refinement protocols in PHENIX were used to refine the structure. Later, a data set was collected from Crystal II, which had the same symmetry as Crystal I, very similar unit-cell dimensions (Table I), and a *d*_*min*_ spacing of 1.75 Å. The Crystal I model was refined against the Crystal II data, with no molecular-replacement step necessary. In the end, individual B-factor, positional, and TLS refinement were used to arrive at the final model, which had excellent geometry, including no Ramachandran outliers (Table I). Riding hydrogen atoms were used at all stages of refinement, and no data were excluded from refinement. The refined structure for Crystal II has been deposited in the Protein Data Bank with accession number 6DET.

### Dynamic Light Scattering

The DLS measurements were taken in a DynaPro NanoStar instrument (Wyatt Technologies, Santa Barbara, CA) in a quartz cuvette containing 5 μL of Tv2483 (2.7 mg/mL) in Sample Buffer at 25 °C. Three technical replicates of three experimental replicates were taken, resulting in nine measurements; for statistical purposes, all replicates were treated equally, and all values presented in the text are unweighted averages ± their respective standard deviations. Reduced intensity autocorrelation functions were exported from Dynamics (Wyatt Technologies) and converted to SEDFIT format with a Python script. The converted reduced field autocorrelation functions were analyzed in SEDFIT with the cumulant model and also with a continuous *R*_*H*_ distribution (11), with a maximum-entropy regularization function at a confidence level of 0.55, and a resolution of 100 between 1 and 500 nm. The viscosity of the buffer was calculated using SEDNTERP (45). Data and fits were plotted in GUSSI (46).

### Analytical Ultracentrifugation

All AUC experiments were conducted in a Beckman XL-I (Indianapolis, IN) centrifuge at 20 °C. Samples (400 μL in Sample Buffer) at three different concentrations (23, 7.3, and 2.3 μM) were introduced into charcoal-filled Epon centerpieces that had been assembled in centrifugation cells such that they had been sandwiched between sapphire windows. The sample cells were placed into a Beckman An50-Ti rotor which was inserted into the centrifuge, placed under vacuum, and incubated at the experimental temperature for 2.5 h. Next, the rotor was accelerated to 50,000 rpm, and data were collected using the absorbance optical system tuned to a wavelength of 280 nm. The data were analyzed using the *c*(*s*) methodology in SEDFIT (47, 48), using maximum-entropy regularization with a confidence level of 0.683, a resolution of 150 between 0 and 10 S, and solution/protein parameters derived from SEDNTERP (45). All distributions were illustrated with and integrated in GUSSI (46), and molar masses were calculated using SEDFIT. All reported values from the SV experiments were the means of these three trials ± the respective standard deviations.

### Differential Scanning Fluorimetry

The thermal stability of Tv2483 and its variants upon ligand binding was performed in a 96-well PCR-plate (Bio-Rad, Hercules, CA) using a Real-Time PCR instrument (Bio-Rad). For ligand screening of physiological compounds (which includes Nitrogen Sources), Biolog’s Phenotype MicroArray (PM) compounds supplied in 96-well microplates (Biolog, Hayward, CA) were dissolved in 50 μL of sterile water to obtain a final concentration of around 10 – 20 mM. Screenings were performed with plate PM3B (Nitrogen Sources). The plate contained 95 compounds and a blank (no ligand) control. The complete plate contents are available in the Biolog website (www.biolog.com).

Each 20 μL standard assay mixture in a 96-well PCR-plate contained 10 μM purified protein and SYPRO Orange (5000X stock solution, Life Technologies, Carlsbad, CA) at 5X concentration in Sample Buffer. Two microliters of the resuspended Biolog compounds were added to each well. Samples were heat denatured from 4 °C to 85 °C in 0.5 °C steps. The protein unfolding curves were monitored from the differential fluorescence changes (-ΔF) of protein-bound SYPRO Orange. The minima of the first derivative values (−*dF*/*dT*) from the raw fluorescence data were used to determine the midpoint of melting temperature (*T*_*m*_).

### Isothermal Titration Calorimetry

All ITC experiments were performed in a Malvern iTC200 (Malvern, UK) at 20 °C. The amino acid under study was always housed in the syringe, while the protein was in the cell (202.9 μL). The stirring speed was 750 rpm. The E128A and R89A proteins were dialyzed against sample buffer. However, the wild-type protein treated thus apparently co-purified with L-arginine and even five days of dialysis were insufficient to remove it, as no heat was observed upon the injection of L-arginine into protein solutions thus treated (not shown). Therefore, we devised a strategy to dislodge the bound ligand. Wild-type Tv2483 (15 mL) was incubated at room temperature overnight in Sample Buffer plus 10 mM L-citrulline. This amino acid was chosen as a competitor because it showed some ability to bind in the DSF assay and because of its structural similarity to L-arginine. The protein was then concentrated to approximately 1 mL in a centrifugal concentrator with a 10,000 Da cutoff (Millipore, Burlington, MA). The protein was then diluted to 15 mL in the same buffer with L-citrulline and incubated for one hour at room temperature. After the same concentration step, the protein was diluted in Sample Buffer (without L-citrulline) to 15 mL. Four more cycles of concentration and dilution were performed. This material could be studied, but still contained a significant amount of material (~35%) that was incompetent to bind to L-arginine or L-citrulline, and it was modeled as such in the analysis. A typical ITC experiment encompassed 19-20 injections of 1.9 to 2.0 μL each, with the first injection being smaller (0.5 to 1.0 μL). For the E128A and R89A proteins, the concentrations in the cell were 100 and 114 μM each, respectively, and the respective syringe concentrations were 1.5 mM and 1.14 mM. For the wild-type protein, a competition study was performed, containing two titrations of L-arginine (500 μM) into Tv2483 (50 μM), two titrations of L-citrulline (250 μM) into Tv2483 (20 μM in one experiment, 50 μM in the other), and one titration of L-arginine (500 μM) into a mixture of L-citrulline and Tv2483 (500 and 50 μM, respectively). All data were integrated using NITPIC (49, 50) and analyzed in SEDPHAT (19, 51). For E128A and R89A, the experiments were conducted in triplicate and the data from each mutant protein were globally analyzed with a 1:1 binding model, allowing the incompetent fraction of the protein to refine. The incompetent fraction for the two mutants were 8% and 12%, respectively. All titrations for the competition experiment with wild-type protein were assembled into a single global analysis using the “A+B+C <-> AB + C <-> AC + B; competing B and C for A” model in SEDPHAT. A single, global correction factor for the incompetent fraction of the protein was applied, and, in the titration where the cell contained L-citrulline, a local concentration-correction factor was also applied. ITC figures were rendered in GUSSI (46).

## Acknowledgments

Results shown in this report are in part derived from work performed at Argonne National Laboratory, Structural Biology Center (SBC) at the Advanced Photon Source. SBC-CAT is operated by UChicago Argonne, LLC, for the U.S. Department of Energy, Office of Biological and Environmental Research under contract DE-AC02-06CH11357.

## FOOTNOTES

Funding was provided by the National Institutes of Health through grant number AI056305 to M.V.N.

The abbreviations used are: AUC, analytical ultracentrifugation; DLS, dynamic light scattering; DSF, differential scanning fluorimetry; GsArtJ, the ArtJ protein from *Geobacillus stearothermophilus*; ITC, isothermal titration calorimetry; LBD, ligand-binding domain; LAO, lysine-arginine-ornithine; StLAO, LAO-binding protein from *Salmonella typhimurium*; r.m.s.d., root-mean-square deviations; SV, sedimentation velocity.

